# Reconstitution of the SARS-CoV-2 ribonucleosome provides insights into genomic RNA packaging and regulation by phosphorylation

**DOI:** 10.1101/2022.05.23.493138

**Authors:** Christopher R. Carlson, Armin N. Adly, Maxine Bi, Yifan Cheng, David O. Morgan

## Abstract

The nucleocapsid (N) protein of coronaviruses is responsible for compaction of the ∼30-kb RNA genome in the ∼100-nm virion. Cryo-electron tomography suggests that each virion contains 35-40 viral ribonucleoprotein (vRNP) complexes, or ribonucleosomes, arrayed along the genome. There is, however, little mechanistic understanding of the vRNP complex. Here, we show that N protein, when combined with viral RNA fragments in vitro, forms cylindrical 15-nm particles similar to the vRNP structures observed within coronavirus virions. These vRNPs form in the presence of stem-loop-containing RNA and depend on regions of N protein that promote protein-RNA and protein-protein interactions. Phosphorylation of N protein in its disordered serine/arginine (SR) region weakens these interactions and disrupts vRNP assembly. We propose that unmodified N binds stem-loop-rich regions in genomic RNA to form compact vRNP complexes within the nucleocapsid, while phosphorylated N maintains uncompacted viral RNA to promote the protein’s transcriptional function.

## Introduction

At different stages of the viral life cycle, viral genomes switch between two distinct structural states: a tightly-packaged protected state inside the virion and a decondensed state that serves as a substrate for translation, transcription, or other processes in the infected cell. The mechanisms that govern the switch between these states are not well understood.

Severe acute respiratory syndrome coronavirus 2 (SARS-CoV-2), the causative agent of the COVID-19 pandemic, is a highly contagious betacoronavirus (*1*). The ∼30-kb single-stranded RNA genome is packed inside the virus in a structure called the nucleocapsid (*2, 3*). Following infection and genomic RNA unpackaging, the first two-thirds of the genome is translated to produce numerous non-structural proteins (Nsps) that rearrange host cell membranes to establish the replication-transcription complex (RTC), a network of double-membrane vesicles that scaffolds viral genome replication and transcription (*4-7*). The final third of the genome then serves as a template for generation of the four structural proteins that form the mature virus (*8-10*).

Transcription of structural protein genes by the viral RNA-dependent RNA polymerase generates negative-sense subgenomic RNAs through a template switching mechanism. These RNAs are then transcribed to positive-sense RNAs, which are translated to produce the Spike (S), Membrane (M), Envelope (E), and Nucleocapsid (N) proteins (*9*). The S, M, and E proteins all contain transmembrane domains that insert into the ER, while the N protein localizes in the cytosol at the RTC and at nearby sites of viral assembly (*4, 6, 11-14*). N protein is the most abundant viral protein in an infected cell (*15*) and serves two essential functions in the coronavirus life cycle: 1) regulation of viral transcription at the RTC, where it facilitates transcriptional template switching required for production of structural protein transcripts; and 2) compaction of the viral RNA genome into the nucleocapsid structure within the virion (*16-18*).

The 46 kDa N protein contains two globular domains flanked by three regions of intrinsic disorder (Fig. 1A) (*19*). The N-terminal domain (NTD) and the C-terminal domain (CTD) bind RNA and are highly conserved among coronaviruses (*20-25*). In solution, N protein exists predominantly as a dimer due to a high-affinity dimerization interface on the CTD but also forms tetramers and higher-order oligomers that are modulated by the disordered N-terminal extension (NTE) and C-terminal extension (CTE, Fig. 1a) (*20, 22, 25-28*). The central disordered region contains a conserved serine/arginine (SR)-rich sequence, which is phosphorylated at multiple sites by host kinases during infection, thereby promoting N protein’s role in viral transcription (*15, 16, 29-31*). The central disordered region also contains sequences that interact with the Nsp3 protein on double-membrane vesicles (*11, 32-34*).

**Figure 1.**
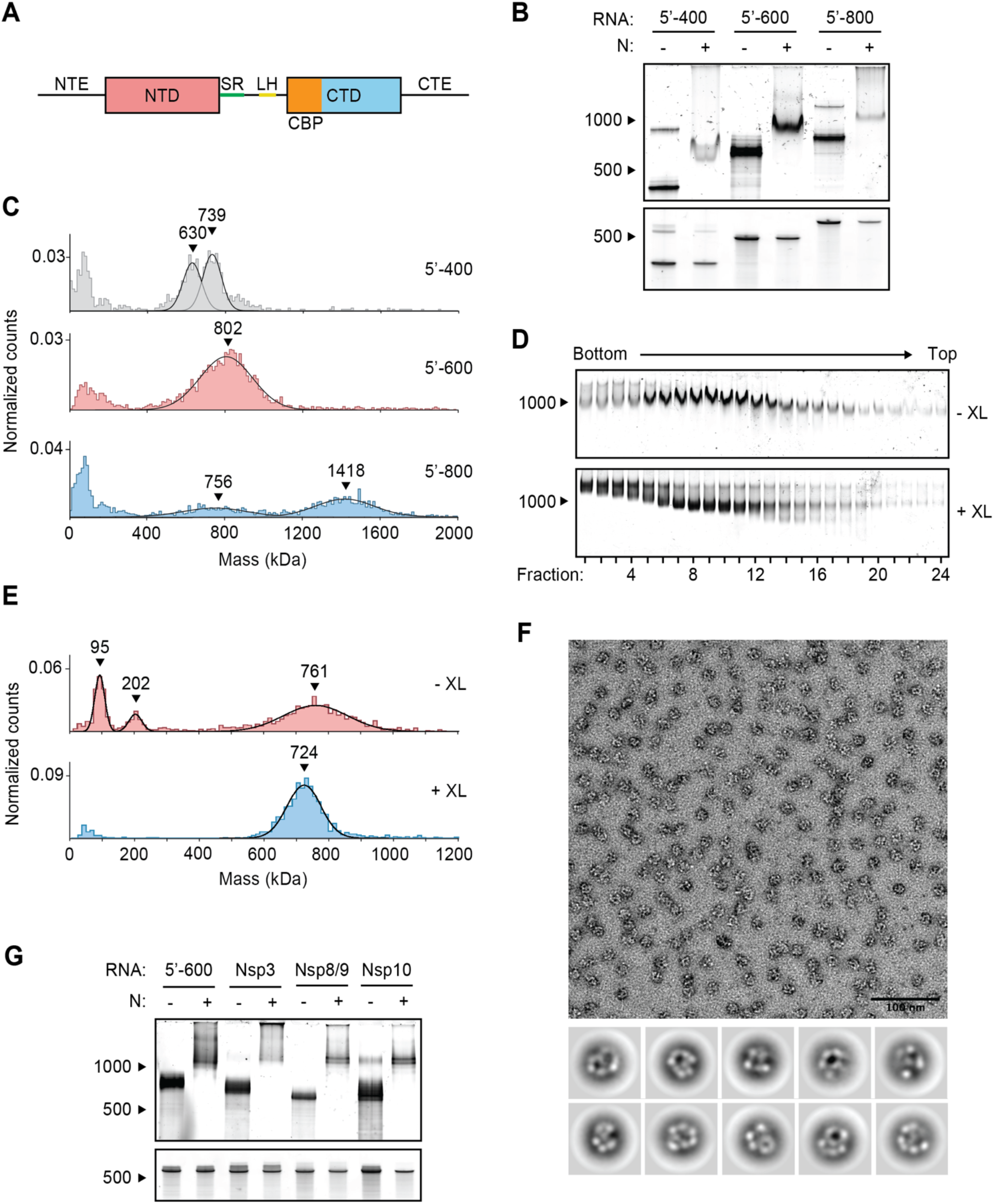
Viral RNA promotes formation of the SARS-CoV-2 ribonucleosome. (**A**) Schematic of N protein domain architecture, including the N-terminal extension (NTE), N-terminal domain (NTD), Serine/Arginine region (SR), Leucine Helix (LH), C-terminal basic patch (CBP), C-terminal domain (CTD), and C-terminal extension (CTE). (**B**) Native (top) and denaturing (bottom) PAGE analysis of 15 μM N protein mixed with 256 ng/μl of the indicated RNA, stained with SYBR Gold to detect RNA species. RNA length standards shown on left (nt). (**C**) Mass photometry analysis of vRNP complexes formed in the presence of 15 μM N and 256 ng/μl RNA. Data were fit to Gaussian distributions, with mean molecular mass indicated above each peak. Representative of two independent experiments (table S1). (**D**) Native gel analysis of glycerol gradient separated vRNP complexes. Top: no crosslinker added (-XL); bottom: 0.1% glutaraldehyde added (+XL) to 40% glycerol buffer (GraFix). (**E**) Fractions 7 and 8 (from D) were combined and analyzed by mass photometry, as in (C). Top: no crosslinker (-XL); bottom: GraFix-purified vRNP (+XL). Representative of two independent experiments (table S1). (**F**) Negative stain electron microscopy and two-dimensional classification of GraFix-purified vRNPs (combined fractions 7 and 8 from D). (**G**) Native (top) and denaturing gel analysis (bottom) of 15 μM N protein mixed with 256 ng/μl of the indicated 600 nt RNA molecules. See table S2 for sequences.

N protein undergoes liquid-liquid phase separation in the presence of viral RNA to form biomolecular condensates (*35-38*). Phosphorylated N combines with RNA to form a liquid-like condensate that might serve as a compartment at the RTC to help protect viral replication and transcriptional machinery from the host cell’s innate immune response (*35, 38, 39*). Unmodified N protein, however, combines with RNA to form more rigid condensates that contain discrete substructures. These gel-like condensates may help to package viral genomic RNA in the nucleocapsid (*35, 37, 38*).

During viral assembly, hypo-phosphorylated N protein binds genomic RNA to form the compact nucleocapsid structure, which is then engulfed by ER membranes containing the S, E, and M proteins to form a mature virus (*4, 5, 16, 30*). Early electron microscopy studies of coronavirus nucleocapsids demonstrated the existence of viral ribonucleoprotein (vRNP) complexes aligned helically along an RNA strand (*40-42*). Recent cryo-electron tomography studies of intact SARS-CoV-2 virions revealed that each virus contains 35-40 discrete, cylindrical nucleosome-like vRNP complexes (*43, 44*). These vRNPs, or ribonucleosomes, are ∼15 nm in diameter and, through low resolution modeling efforts, are speculated to contain twelve N proteins in complex with up to 800 nt of RNA (30,000 nt ÷ 38 vRNPs = 800 nt). A ‘beads-on-a-string’ model has been proposed as a general mechanism of coronavirus packaging: vRNPs (the beads) locally compact RNA within the long genomic RNA strand (the string).

Unlike string, however, the SARS-CoV-2 genomic RNA is highly structured, containing an elaborate array of heterogeneous secondary and tertiary structural elements that are present in both infected cells and in the virion (*45, 46*). Thus, N protein must accommodate a variety of RNA structural elements to form the compact vRNPs of the nucleocapsid. Mechanistic insight into this model and overall vRNP architecture is lacking.

In our previous work, we observed that purified N protein and a 400-nt viral RNA fragment assemble into vRNP particles similar to those seen inside the intact virus, suggesting that N protein and RNA alone are sufficient to form the vRNP (*38*). Here, we explore the biochemical properties, composition, and regulation of these particles. We find that vRNPs form in the presence of stem-loop-containing RNA though a multitude of protein-protein and protein-RNA interactions. Phosphorylation of N protein weakens these interactions and inhibits the formation of ribonucleosomes.

## Results

### Stem-loop-containing RNA promotes ribonucleosome formation

We previously observed vRNP complexes in vitro when N protein was mixed with a 400-nt viral RNA from the 5’ end of the genome, while cryo-electron tomography studies of intact viruses suggest that the vRNP packages up to 800 nt of RNA (*38, 43, 44*). To further investigate the impact of RNA length on vRNP assembly, we mixed N protein with 400-, 600-, and 800-nt RNA fragments from the 5’ end of the genome (5’-400, 5’-600, and 5’-800, respectively) and analyzed the resulting complexes by electrophoresis on a native TBE gel. All three RNAs shifted to a larger species in the presence of N (Fig. 1B), indicating that N protein bound the RNAs and retarded their electrophoretic mobility. N protein in complex with 5’-600 RNA resulted in a particularly discrete, intense band, suggesting that it forms a stable RNA-N protein complex.

We used mass photometry to better characterize these RNA-N protein complexes. Mass photometry uses light scattering to measure the mass of single molecules in solution, resulting in a histogram of mass measurements centered around the average molecular mass of the protein complex. N protein in complex with 5’-400 RNA resulted in two mass peaks that were smaller than the single broad peak of N protein bound to 5’-600 RNA, suggesting the 5’-400 vRNP was not fully assembled and contained subcomplexes (Fig. 1C). N protein mixed with 5’-800 RNA formed two broad peaks: one smaller peak that appears similar in size to the 5’-600 species (both ∼750-800 kDa), and a second larger peak roughly twice as large as the first (∼1400 kDa). This suggests that one (∼750 kDa) or two vRNPs (∼1400 kDa) can form on a single 5’-800 RNA molecule (Fig. 1C). The 5’-600 RNA was therefore chosen as a representative viral RNA to further study the ribonucleosome.

To purify the vRNP complex for more detailed analysis, N protein was mixed with 5’-600 RNA and separated by centrifugation on a 10-40% glycerol gradient. Individual fractions were analyzed by native gel electrophoresis (Fig. 1D, top), after which fractions 7 and 8 were combined for analysis by mass photometry. We observed three major peaks centered at 97 ± 2 kDa, 207 ± 6 kDa, and 766 ± 6 kDa (Fig. 1E, top and table S1). These peaks likely correspond to free N protein dimer (predicted mass 91.2 kDa; see Fig. 2D, top), unbound 5’-600 RNA (predicted mass 192.5 kDa), and the vRNP complex, respectively. The presence of free N protein dimer and unbound RNA suggests that the vRNP complex dissociated upon dilution for mass photometry analysis.

**Figure 2.**
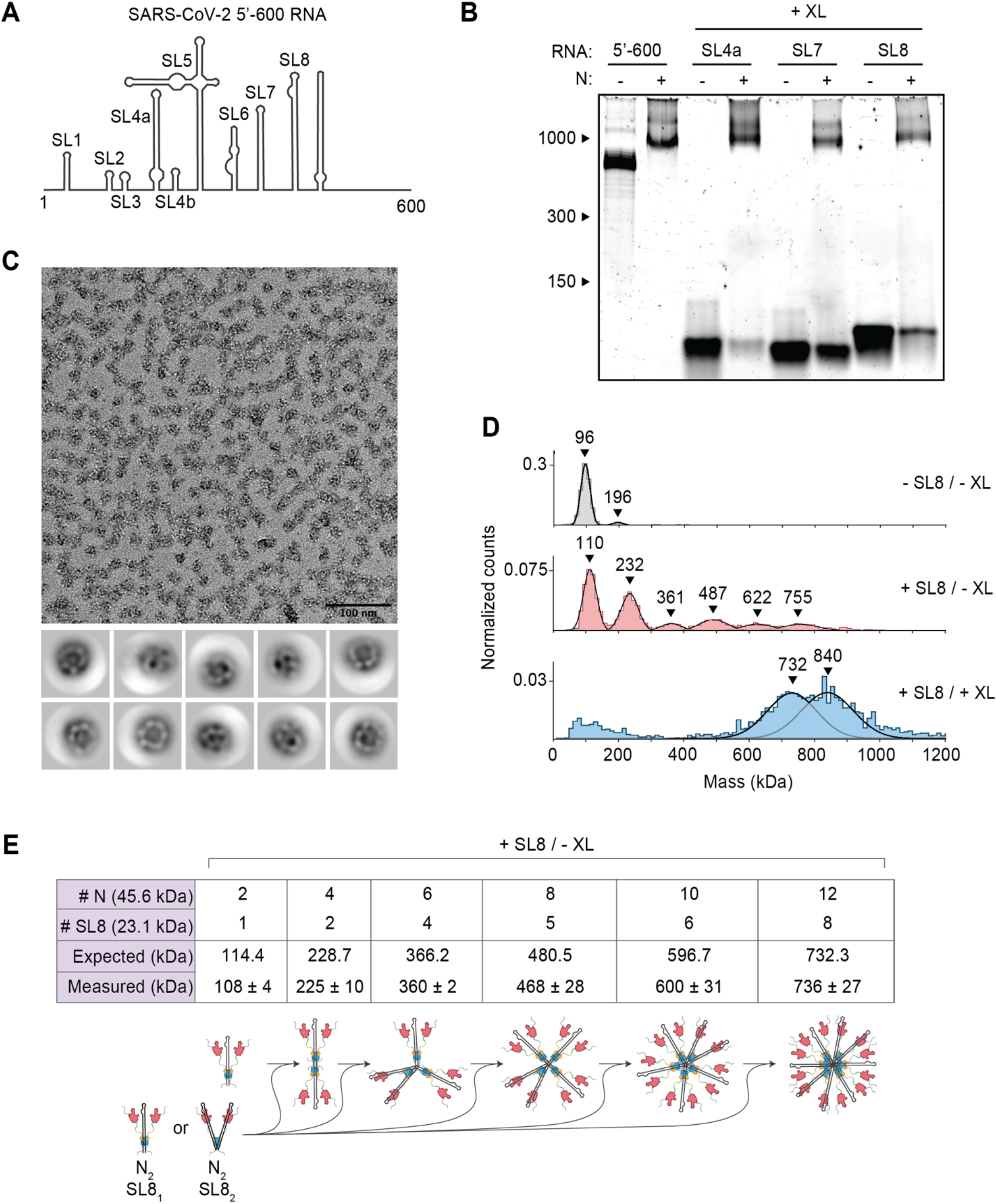
Stem-loop RNA, in complex with N protein, drives ribonucleosome formation. (**A**) Schematic of RNA secondary structure in the 5’-600 RNA (*45*). (**B**) Native gel analysis of 15 μM N protein mixed with 256 ng/μl of the indicated RNAs. Samples containing stem-loop RNA were crosslinked to stabilize the resulting complex, while the 5’-600 RNA sample was left un-crosslinked. Corresponding denaturing gel analysis shown in fig. S1A. (**C**) Fractions 7 and 8 of GraFix-purified SL8 assembled vRNPs were combined and analyzed by negative stain electron microscopy and two-dimensional classification. (**D**) Mass photometry analysis of indicated N protein-RNA mixtures. Top: N protein alone; middle: N protein mixed with SL8, un-crosslinked; bottom: crosslinked complexes of N protein bound to SL8 (data reproduced from fig. S1B for ease of comparison). Representative of two independent experiments (table S1). (**E**) Predictions of N protein and RNA stoichiometry, based on measured masses of N protein in complex with SL8 RNA without crosslinker (D, middle panel). Measured masses are means ± standard deviation in two independent experiments (table S1). Below the table is a schematic of a proposed assembly mechanism in which N protein dimers, bound to one or two stem-loop RNAs, iteratively assemble to the full vRNP.

To stabilize the complex, a crosslinker (0.1% glutaraldehyde) was added to the 40% glycerol buffer, creating a gradient of glutaraldehyde throughout the glycerol to crosslink the protein complex during centrifugation (a technique known as gradient fixation, or GraFix) (*47*). Analysis of the GraFix-purified fractions by native gel electrophoresis revealed sharper, more discrete bands compared to the non-crosslinked sample (Fig. 1D, bottom). The distribution of vRNP complexes across the gradient was similar between the two conditions. The GraFix purified sample (fractions 7 + 8) was analyzed by mass photometry, revealing one peak with an approximate mass of 727 ± 4 kDa (Fig. 1E, bottom and table S1). This is consistent with the idea that the non-crosslinked sample dissociates upon dilution for mass photometry and suggests a likely stoichiometry of 12 N proteins (547.5 kDa) bound to one 5’-600 RNA (192.5 kDa; total predicted mass: 740 kDa). Alternatively, a complex of 8 N proteins (365 kDa) with two 5’-600 RNAs (385 kDa; total predicted mass: 750 kDa) is also consistent with these results.

Negative stain electron microscopy (EM) of the GraFix-purified sample revealed discrete 15-nm particles with an electron-dense center surrounded by an outer ring (Fig. 1F). Two-dimensional classification revealed particles with variable composition and conformation, suggesting inherent structural heterogeneity within the vRNP complex that may reflect the diverse RNA stem-loop structures in the 600 nt RNA strand (see Fig. 2A). While these averages are heterogeneous, they are similar in size and shape to vRNP complexes previously observed within SARS-CoV-2 virions by cryo-electron tomography (*43, 44*).

We next tested if specific RNA sequences or regions of the genome promote formation of the vRNP. Four 600-nt genomic regions were transcribed in vitro, individually mixed with N protein, and analyzed by native gel electrophoresis: (1) 5’-600 (nucleotides 1–600), (2) Nsp3 (nucleotides 7,800–8,400), (3) Nsp8/9 (nucleotides 12,250–12,850), (4) Nsp10 (nucleotides 13,200–13,800). All RNAs appeared to form appropriately-sized vRNPs, although the Nsp3 RNA appeared less effective (Fig. 1G). These results suggest that the vRNP can accommodate a variety of viral RNA and does not require specific sequences to form, although certain sequences may form more stable ribonucleosomes.

We then sought to explore the relationship between RNA structure and vRNP formation by dissecting the structures required for vRNP assembly within the 5’-600 RNA. This highly structured 600 nt genomic region contains several well-characterized stem-loops varying in size from 20 to ∼150 nt (Fig. 2A) (*45, 48*). Three stem-loop RNAs (SL4a, 56 nt; SL7, 46 nt; SL8, 72 nt) were individually mixed with N protein, crosslinked (with 0.1% glutaraldehyde) to stabilize the resulting complexes and assessed for vRNP formation by native gel electrophoresis. Appropriately sized vRNP complexes formed in the presence of all three stem-loops (Fig. 2B and fig. S1A). Each crosslinked complex was analyzed by mass photometry. SL4a mixtures contained three broad peaks of 515 ± 19 kDa, 739 ± 5 kDa, and 876 ± 1 kDa (fig. S1B, top, and table S1). SL7 also generated three mass peaks at 502 ± 26 kDa, 615 ± 22 kDa, and 713 ± 13 kDa (fig. S1B, middle, and table S1). SL8 generated two peaks at 737 ± 11 kDa and 840 ± 12 kDa (fig. S1B, bottom, and table S1) and was chosen for further analysis due to less heterogeneity in the composition of the complex.

SL8-containing vRNPs were purified by GraFix (fig. S1C). Analysis of peak fractions (7 + 8) by mass photometry (fig. S1D) indicated that SL8 vRNPs were similar in mass to vRNPs assembled with 5’-600 RNA (Fig. 1E). Negative stain EM (Fig. 2C) and two-dimensional class averages revealed ring structures that resemble vRNPs assembled with the 5’-600 RNA (Fig. 1F).

These data suggest that ribonucleosome formation does not require 600 continuous bases of RNA but can be achieved with multiple copies of a relatively short and simple stem-loop structure. Unlike the 5’-600 RNA, the short stem-loop RNA is unlikely to serve as a platform to recruit multiple copies of N protein to assemble a vRNP. We speculate that the binding of a stem-loop RNA to N protein induces a conformational change that promotes protein-protein interactions required for vRNP formation. In the more physiologically relevant context of long RNAs, these weak protein-protein interactions are likely stabilized by multivalent interactions with an RNA molecule.

To test the requirement for secondary structure in vRNP formation, we analyzed a mutant SL8 (mSL8) carrying 12 mutations predicted to abolish the stem-loop structure. vRNP formation was reduced in the presence of mSL8, suggesting that ribonucleosomes form more readily with double-stranded stem-loop structures (fig. S1E).

Analysis of non-crosslinked SL8-N protein complexes shed further light on vRNP assembly. Mass photometry of the SL8-N sample revealed a major species at ∼110 kDa, with five evenly spaced complexes every 120-130 kDa thereafter up to ∼755 kDa (108 ± 4 kDa, 225 ± 10 kDa, 360 ± 2 kDa, 468 ± 28 kDa, 600 ± 31 kDa, 736 ± 27 kDa) (Fig. 2D, middle, and table S1). N protein alone exists primarily as a ∼96 kDa dimer (97 ± 1 kDa) at the low concentration used for mass photometry (Fig. 2D, top, and table S1; predicted mass 91.2 kDa), so the ∼110 kDa peak likely represents one N protein dimer bound to one SL8 RNA (predicted mass of RNA: 23.1 kDa; predicted mass of complex: 114.4 kDa). The stepwise ∼120-130 kDa increases in molecular mass are consistent with the addition of an N dimer bound to either one or two SL8 RNA molecules (predicted mass: 114.4 kDa or 137.5 kDa, respectively). These results support a potential assembly mechanism in which N protein dimers, bound to one or two stem-loops, iteratively assemble to form a full ribonucleosome containing twelve N proteins and six to twelve stem-loop RNAs (Fig. 2E). These data support the possibility that the vRNP assembled with 5’-600 RNA (Fig. 1E) contains 12 N proteins bound to one RNA.

In some crosslinked vRNP preparations, we observed an additional large peak in mass photometry that is likely to contain more than 12 N proteins. As mentioned above, crosslinked SL8 vRNPs contain a broad peak of 840 ± 12 kDa in addition to the 737 ± 11 kDa peak (fig. S1B, bottom; also shown in Fig. 2D, bottom). Based on the similar molecular mass of the smaller peak in the crosslinked sample (737 ± 11 kDa) to the non-crosslinked sample (736 ± 27 kDa), we suspect that the larger crosslinked complex of 840 ± 12 kDa contains 14 N proteins. These results suggest that the ribonucleosome defaults to a stable complex of 12 N proteins bound to a variable number of RNA stem-loops but can adapt to accommodate fewer or more N protein dimers bound to additional RNA.

### Multiple N protein regions promote formation of the ribonucleosome

Next, we sought to explore the regions of the N protein required for vRNP formation. We analyzed mutant proteins lacking the following regions: (1) the 44-aa N-terminal extension (NTE), a poorly conserved prion-like sequence that promotes RNA-induced liquid-liquid phase separation of N protein (*38, 49*); (2) the highly conserved 31-aa serine/arginine (SR) region that has been implicated in RNA binding, oligomerization, and phosphorylation (*15, 16, 29-31, 50-52*); (3) the 20-aa leucine helix (LH), an alpha helix downstream of the SR region that interacts with Nsp3 (*33*); (4) the 33-aa CTD basic patch (CBP), which forms a highly basic RNA-binding groove on the CTD and has been implicated in helical stacking of N protein (*22, 53*); and (5) the 55-aa C-terminal extension (CTE), which has been implicated in tetramerization and oligomerization of N (*20, 50, 52, 54*) (Fig. 3A and fig. S2A).

**Figure 3.**
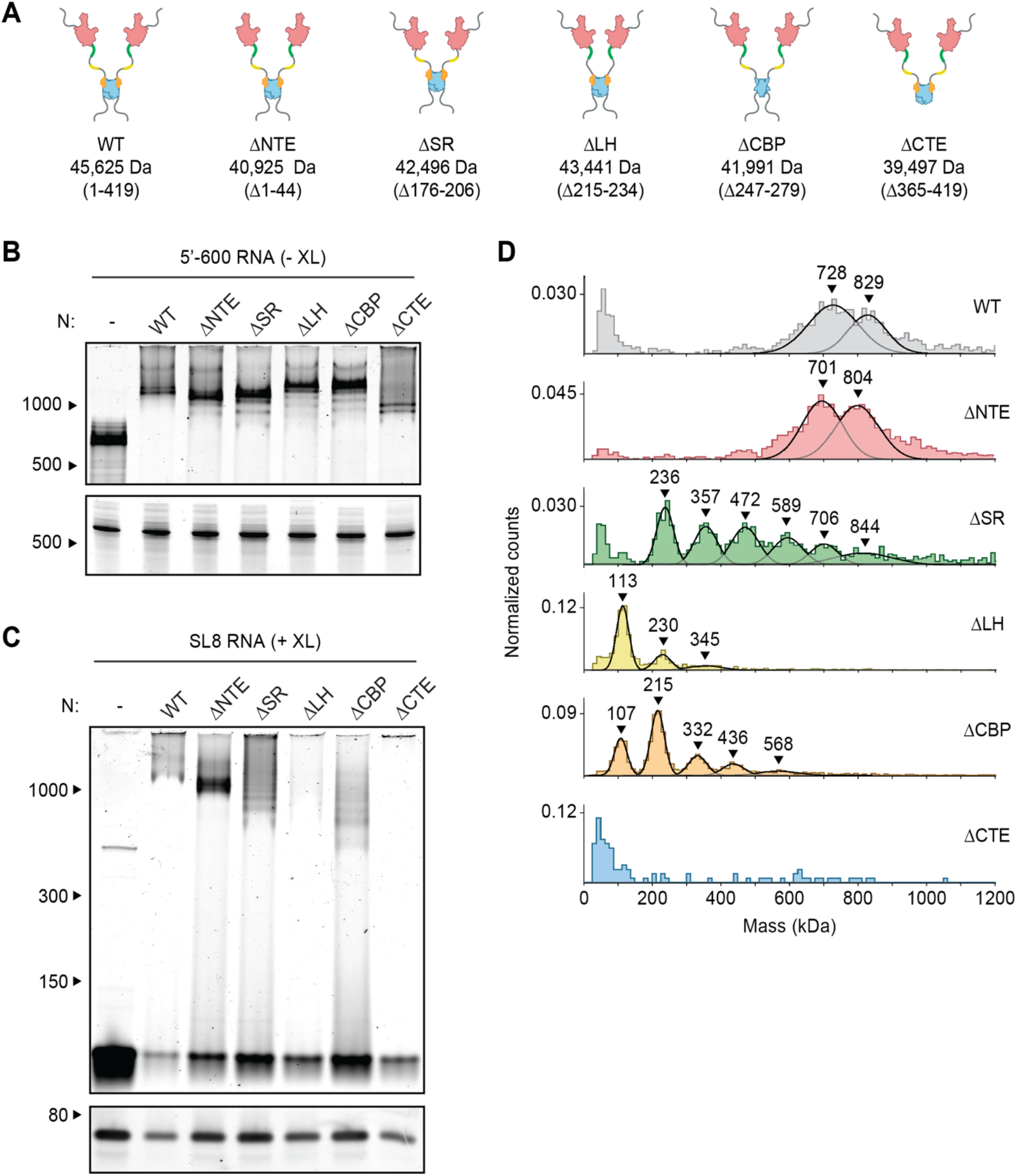
Disordered regions contribute to vRNP formation. (**A**) Schematic of wild-type (WT) N protein and deletion mutants, as described in the text. Mass is that of monomeric N protein. (**B**) 15 μM N protein mutants were mixed with 256 ng/μl 5’-600 RNA and analyzed by native (top) and denaturing (bottom) gel electrophoresis. (**C**) 20 μM N protein mutants were mixed with 256 ng/μl SL8 RNA and analyzed by native (top) and denaturing (bottom) gel electrophoresis. SL8 ribonucleoprotein complexes were crosslinked prior to analysis. (**D**) Mass photometry analysis of crosslinked N protein mutants (20 μM) bound to SL8 RNA (256 ng/μl). Representative of at least two independent experiments (table S1).

Mutant N proteins were mixed with 5’-600 RNA and analyzed by native gel electrophoresis (Fig. 3B). All mutant N proteins, with the exception of the CTE deletion, appeared to form fully assembled vRNPs. Most mutants contained varying amounts of lower bands beneath the fully shifted vRNP. These lower bands might represent sub-complexes in which the 5’-600 RNA is bound to fewer N proteins, presumably due to defects in vRNP assembly or stability. Deletion of the CTE resulted in a small shift that was considerably lower than the fully shifted vRNP. These results suggest that the ΔCTE N protein binds RNA but fails to form the fully assembled vRNP, hinting at an important role for the CTE in vRNP formation.

Studies of deletion mutants in complex with SL8 RNA, which minimizes the contribution of multivalent RNA binding, allowed us to investigate the critical protein-protein interactions that contribute to ribonucleosome formation. Mutant N proteins were mixed with SL8 RNA, crosslinked, and analyzed by native gel electrophoresis and mass photometry (Fig. 3C, D, and table S1). Deletion of the NTE had little effect, other than to decrease the size of the vRNP complexes by ∼30-40 kDa, suggesting that the NTE is not required for vRNP formation. All other deletion mutants had major defects in vRNP assembly.

Deletion of the CTE and LH resulted in almost complete disappearance of the vRNP when analyzed by native gel electrophoresis (Fig. 3C). These mutants appeared cloudy, and turbidity analysis revealed a higher absorbance at 340 nm compared to wild-type, suggesting formation of biomolecular condensates (fig. S2B). Mass photometry analysis of the LH deletion showed a dominant peak at ∼110 kDa, with two minor peaks at ∼230 kDa and ∼345 kDa (Fig. 3D and table S1). The smallest peak represents an N dimer bound to one SL8 RNA, with the next two representing stepwise additions of one or two N dimers bound to an RNA. Thus, protein-protein interactions mediated by the LH are required for vRNP formation. Deletion of the CTE resulted in no discernable peaks above background on the mass photometer (Fig. 3D), further confirming the essential role of the CTE in vRNP formation and suggesting that tetramerization driven by the CTE is required for ribonucleosome formation or stability.

Deletion of the SR and CBP regions also resulted in defects in vRNP assembly; both mutants exhibited a laddering of ribonucleoprotein subcomplexes when analyzed by native gel electrophoresis, as well as stepwise 120-130 kDa increases in molecular mass revealed by mass photometry (Fig. 3C, D, and table S1). These data suggest the SR and CBP regions are required for complete assembly of the ribonucleosome.

LH and CBP deletions resulted in a minimal ribonucleoprotein complex of ∼110 kDa, consistent with one N protein dimer bound to one SL8 RNA molecule. Interestingly, the SR deletion resulted in a minimal ribonucleoprotein complex of ∼230 kDa, consistent with one N protein tetramer bound to two SL8 RNAs.

Native gel analysis revealed an increase in free SL8 RNA in the various mutant N protein samples compared to wild-type, suggesting defects in RNA binding (Fig. 3C). We performed fluorescence anisotropy to quantitatively measure the affinity of a 10-nt RNA (of random sequence) for mutant N proteins, which likely reflects RNA binding to the high-affinity RNA-binding site on the NTD (fig. S2C) (*51*). Wild-type N protein had an affinity of 28 ± 6 nM for the 10 nt RNA oligo, which is consistent with previous measurements of RNA binding to the NTD (*51*). All mutant N proteins, except the SR deletion, had similar affinities for RNA. Deletion of the SR region resulted in a modest ∼5-fold decrease in affinity, consistent with previous reports that the SR region makes a slight contribution to RNA binding at the NTD (*51*). The increase in free SL8 RNA in native gel electrophoresis is therefore likely caused by defects in lower-affinity RNA binding at other sites in the N protein, which might be necessary for proper ribonucleosome formation.

### Phosphorylation inhibits formation of the ribonucleosome

The SR region of N protein is heavily phosphorylated in cells infected by SARS-CoV-2, and this modification is required for the protein’s role in viral transcription (*15, 16, 29-31*). In contrast, N protein in the virion is thought to be poorly phosphorylated (*16, 30*). We previously observed defects in vRNP formation when the 5’-400 RNA was mixed with a phosphomimetic N protein (the 10D mutant, in which 10 serines and threonines in the SR region are replaced with aspartic acid) (*38*), and here we sought to further explore phosphoregulation of the ribonucleosome. We mixed 5’-600 RNA with the 10D mutant and analyzed vRNP formation by native gel electrophoresis (Fig. 4A, left). The mutant formed an appropriately sized vRNP, apart from minor subcomplexes formed below the fully assembled vRNP. GraFix purification of the 10D mutant in complex with 5’-600 RNA revealed a range of ribonucleoprotein complexes similar to that seen with wild-type N protein (Fig. 4B and Fig. 1D, bottom). Mass photometry of fractions 7 + 8 confirmed a similar mass of the 10D and wild-type vRNPs, apart from a minor ∼830 kDa peak observed with the 10D mutant (fig. S3A and Fig. 1E, bottom).

**Figure 4.**
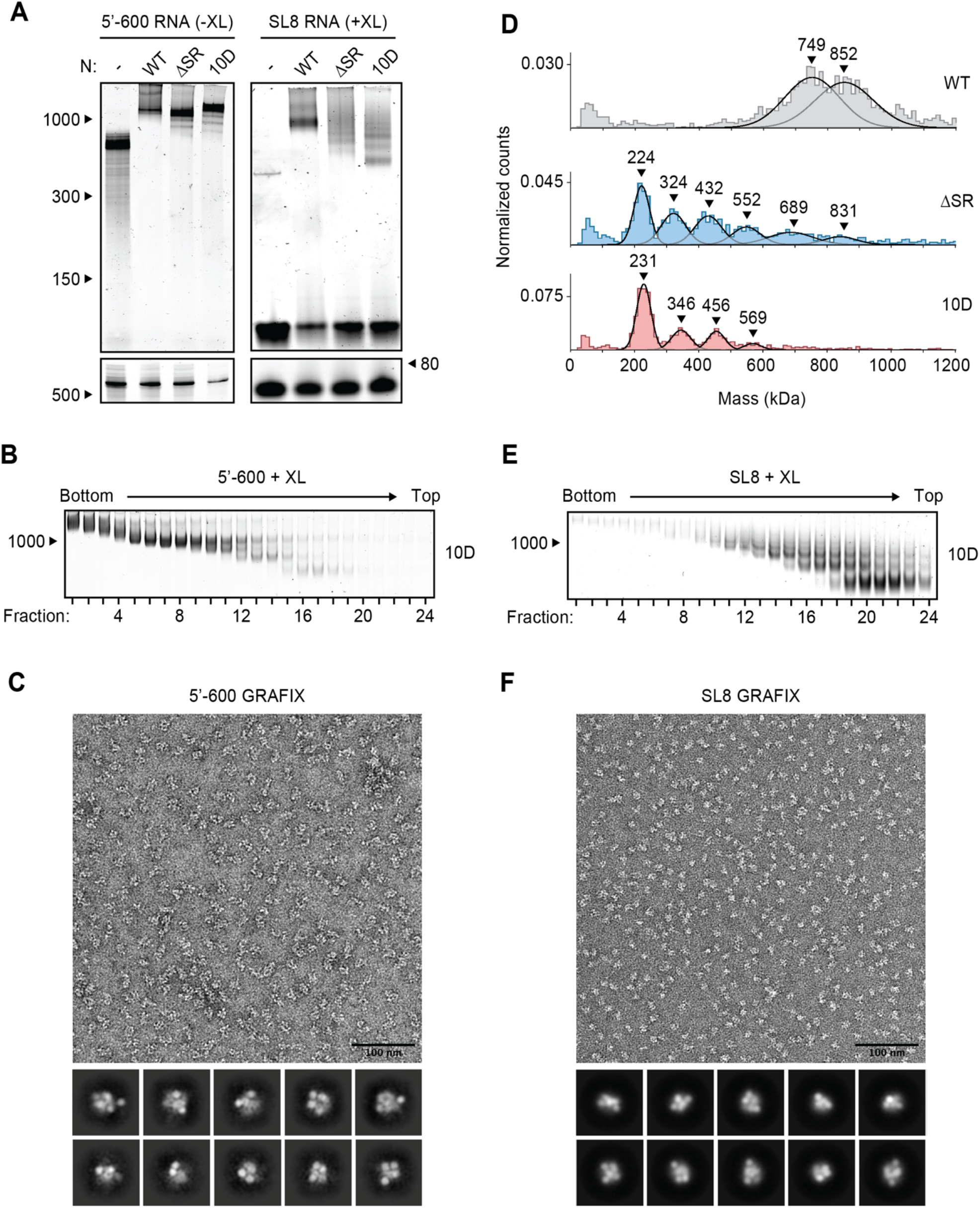
Phosphomimetic mutations in the SR region of N prevent vRNP assembly. (**A**) 15 μM N protein constructs were combined with 256 ng/μl 5’-600 RNA (left) or 256 ng/μl SL8 RNA (right) and analyzed by native (top) and denaturing (bottom) gel electrophoresis. SL8 ribonucleoprotein complexes were crosslinked prior to native gel electrophoresis. WT, wild type. See Fig. 5A for the ten sites of phosphorylation mutated to aspartic acid in the 10D mutant. (**B**) 15 μM phosphomimetic N protein (10D) was mixed with 256 ng/μl 5’-600 RNA and separated by glycerol gradient centrifugation in the presence of crosslinker (GraFix). Fractions were collected and analyzed by native gel electrophoresis. (**C**) Fractions 7 and 8 of GraFix-separated vRNPs (from B) were combined and analyzed by negative stain electron microscopy and two-dimensional classification. (**D**) 15 μM N protein mutants were mixed with 256 ng/μl SL8 RNA, crosslinked, and analyzed by mass photometry. Representative of at least two independent experiments (table S1). A separate analysis of ΔSR mutant is also shown in Fig. 3D. (**E**) 15 μM 10D N protein was mixed with 256 ng/μl SL8 RNA and separated by GraFix. Fractions were analyzed by native gel electrophoresis. (**F**) Fractions 19 and 20 of GraFix-purified 10D N in complex with SL8 RNA (from E) were combined and visualized by negative stain electron microscopy and two-dimensional classification.

Negative stain EM and two-dimensional class average analysis of the GraFix-purified 10D ribonucleoprotein complex, however, revealed a markedly different structure compared to the wild-type vRNPs (Fig. 4C and Fig. 1F). The 10D complex appears extended and heterogeneous, unlike the compact structure of the wild-type vRNP, and does not average into discrete, recognizable two-dimensional classifications. We therefore speculate that the 600 nt RNA provides sufficient binding sites for twelve 10D N proteins, but the 10D mutant is unable to condense into the ring structure observed with the wild-type N protein. We note that there was no defect in RNA binding to the NTD of the 10D mutant (K_d_ = 31 ± 7 nM) (fig. S2C).

Ribonucleosome formation by the 10D mutant with the SL8 RNA was severely reduced when analyzed by native gel electrophoresis and mass photometry (Fig. 4A, right, and Fig. 4D). Both assays revealed a laddering of vRNP complexes, consistent with an inability of the 10D mutant to form a stable, fully assembled vRNP. Purification of the 10D + SL8 complex by GraFix revealed a clear shift toward lower molecular mass species when compared to wild-type N (Fig. 4E compared to fig. S1C). This result was confirmed by mass photometry analysis of fractions 19 + 20 (fig. S3B). Interestingly, the minimal unit of vRNP complex assembly with the 10D mutant (like the SR deletion) is ∼230 kDa, which is consistent with an N protein tetramer bound to two SL8 RNAs. Negative stain EM and two-dimensional class averages of the GraFix-purified complex (fractions 19 + 20) revealed a smaller overall structure with an electron density distribution clearly distinct from vRNP complexes formed by wild-type N (Fig. 4F compared to Fig. 2C).

We next tested vRNP assembly with N protein that had been phosphorylated in vitro. In recent work, Yaron et al. (*29*) elegantly demonstrated a multi-kinase cascade that results in maximally phosphorylated N protein: SRPK phosphorylates S188 and S206, which primes the protein for subsequent phosphorylation of 8 more sites within the SR by GSK3, which then primes a final 4 sites for phosphorylation by CK1 (Fig. 5A). Consistent with this model, we observed maximal phosphorylation of N in the presence of all three kinases (Fig. 5B). Phosphorylation was greatly reduced when both SRPK priming sites were mutated to alanine (S188A + S206A mutant) (Fig. 5B). We mixed kinase-treated wild-type or S188A + S206A N proteins with SL8 RNA and purified the resulting vRNP complexes by GraFix (Fig. 5C). Wild-type phosphorylated N protein migrated as a low molecular weight ribonucleoprotein complex across the gradient, similar to the 10D mutant. The poorly phosphorylated S188A + S206A mutant, however, formed an appropriately sized vRNP across the gradient, similar to wild type unphosphorylated N protein (Fig. 5C). Mass photometry of the GraFix-purified samples further substantiated the defect in wild-type phospho-N vRNP assembly (Fig. 5D, top), which is rescued by mutation of the two priming phosphorylation sites (the S188A + S206A mutant) (Fig. 5D, bottom).

**Figure 5.**
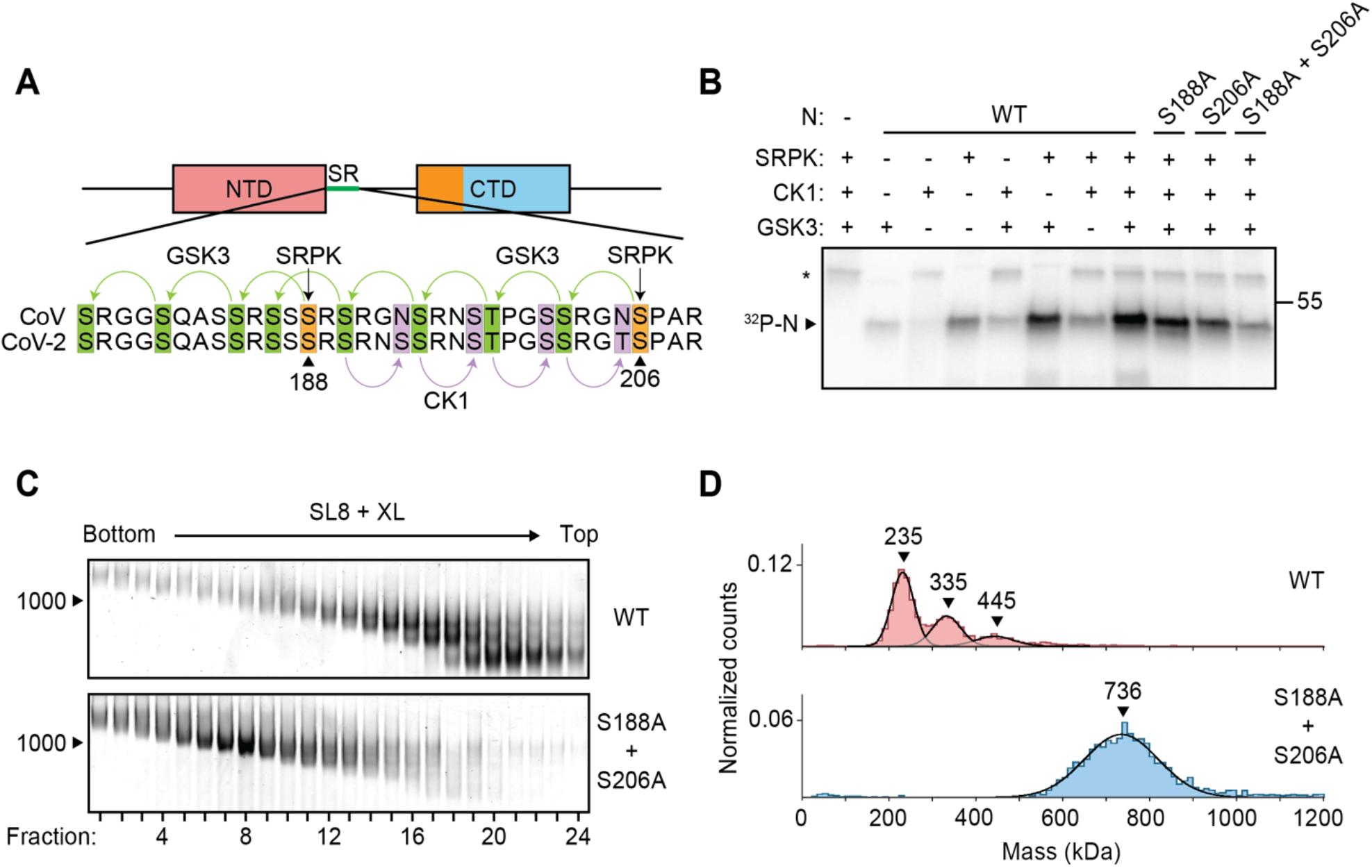
Phosphorylation of N protein inhibits ribonucleosome formation. (**A**) Sequence of N protein SR regions from SARS-CoV (aa 177-210) and SARS-CoV-2 (aa 176-209). The proposed mechanism of sequential phosphorylation (*29*) is initiated by SRPK at S188 and S206 (orange), which leads to downstream phosphorylation of eight sites by GSK3 (green), allowing for final phosphorylation of four additional sites by CK1 (purple). In the phosphomimetic 10D mutant used in Fig. 4, the SRPK and GSK3 sites are changed to aspartic acid. (**B**) Wild-type (WT) and mutant N protein constructs were incubated with the indicated kinases in the presence of radiolabeled ATP and analyzed by SDS-PAGE and autoradiography. Phosphorylated N is indicated. Asterisk denotes autophosphorylation of CK1. Molecular mass marker shown on right (kDa). (**C**) N protein (WT or S188A + S206A) was phosphorylated by SRPK, GSK3, and CK1, and then mixed with SL8 RNA. The resulting ribonucleoprotein complexes were separated by glycerol gradient centrifugation in the presence of crosslinker (GraFix) and analyzed by native gel electrophoresis. (**D**) Peak fractions from the GraFix analyses in C were analyzed by mass photometry. Top: fractions 19 + 20 of wild-type N; bottom: fractions 7 + 8 of S188A + S206A mutant N. Representative of two independent experiments (table S1).

## Discussion

The ‘beads-on-a-string’ model for coronavirus genome packaging lacks mechanistic detail. Here, we demonstrate that the N protein of SARS-CoV-2 assembles with viral RNA in vitro to form ribonucleosomes. These structures, which have been observed previously in intact SARS-CoV-2 virions by cryo-electron microscopy (*43, 44*), likely contain twelve N proteins (6 dimers) and a variable number of stem-loop RNA structures.

Short stem-loop RNAs appear to induce conformational changes in N protein that promote protein-protein interactions necessary for ribonucleosome assembly. These interactions might involve SR binding to the CTD (*52*), LH binding to other regions of the N protein, helical stacking of the CBP (*22*), and tetramerization driven by the CTE (*20, 50, 54, 55*). All of these binding interfaces contribute to the stability of the vRNP, but the CTE seems particularly critical for ribonucleosome formation.

vRNPs formed with long viral RNA (600 nt) do not fall apart as readily when diluted for mass photometry and do not require crosslinking for visualization by native gel electrophoresis. These results suggest that vRNPs assembled with 600-nt RNAs are more stable than those formed with multiple copies of a single short stem-loop RNA, potentially because a single long RNA provides binding sites for all twelve N proteins in the vRNP. This multivalent RNA scaffold stabilizes low-affinity protein-protein interactions within the vRNP and reflects the more physiologically relevant state of RNA compaction by N protein in the virion.

Phosphorylation of N protein in its disordered SR region by host kinases inhibits vRNP formation, perhaps by inhibiting SR-dependent protein-protein interactions, lending further mechanistic insight into the functions of N protein phosphorylation and dephosphorylation during coronavirus infection. Despite the high level of N protein phosphorylation in the infected cell, mechanisms must exist to generate a poorly phosphorylated N protein population at sites of viral assembly.

Coronavirus genomic RNA is structurally heterogeneous (*45, 46*), and it remains unclear how ribonucleosomes accommodate variable RNA sequence and structure to package RNA in the virion. We find that the vRNP assembles in the presence of 600-nt RNA fragments from multiple genomic regions, suggesting that no specific sequences are required for vRNP formation. Furthermore, the ability of short stem-loop RNAs to trigger vRNP formation suggests that ribonucleosome formation does not require 600 continuous bases of RNA. Inside the virion, it is not known whether each ribonucleosome forms on a continuous stretch of RNA in a nucleosome-like fashion or instead acts as a hub that binds stem-loops distributed across the genome, creating a web of condensed, interlinked protein-RNA interactions with ‘nodes’ at the ∼38 vRNPs.

Studies of ribonucleosome assembly with a small stem-loop RNA demonstrate that the vRNP is compositionally adaptive – that is, it can contain a variable number of N protein dimers bound to a variable number of stem-loop RNAs and assembles by iterative additions of N protein dimers bound to stem-loop RNAs. Our data suggest that the most stable form of the vRNP is 12 N proteins in complex with ∼600 nt of RNA, but we also observed complexes that contain fewer or more N protein dimers. Given the iterative assembly of the vRNP, the multitude of protein-protein and protein-RNA interactions, and the high concentrations of N protein and RNA in the nucleocapsid, it seems reasonable to expect that the vRNP can expand to expose binding sites that allow additional N protein dimers to insert themselves into, or dissociate from, the cylindrical vRNP complex.

Our results, together with data from other studies, provide insights into the general architecture of coronavirus RNA packaging. There are ∼38 vRNPs per virus (*43, 44*), with each vRNP likely containing ∼12 N proteins in complex with ∼600 bases of viral RNA. This suggests that within a virus, the vRNPs contain ∼500 N proteins bound to ∼23,000 nt of RNA. It has been estimated that there are ∼1000 N proteins per virus (*56*), while the viral genome is 30,000 nt in length, suggesting that some N proteins and RNA in the virion are not incorporated into vRNPs. Cryo-electron tomography studies indicate that most vRNPs are associated with the inner face of the membrane envelope, with a structure-free center in every virus (*40, 43, 57*). Based on previous studies from our lab and others, this central region in the virion might contain a gel-like condensate of N protein bound heterogeneously to viral RNA (*35, 37, 38*).

During viral assembly, one copy of the ∼30 kb viral genome is packaged per virus, while cellular and subgenomic viral RNA are excluded from the virion (*58*). In murine hepatitis virus (MHV), a 94 nt stem-loop in the genomic RNA is necessary for exclusion of subgenomic RNA from the virus, suggesting that an analogous sequence or structure exists in SARS-CoV-2 (*59*). Our results demonstrate that the vRNP of SARS-CoV-2 does not appear to possess strict sequence or structure specificities, suggesting that another mechanism ensures specific incorporation of genomic RNA into the mature virus. The Membrane (M) protein likely functions in this capacity.

M protein is a 25 kDa structural protein containing three transmembrane helices followed by a ∼100 aa C-terminal domain that faces the interior of the virion and is thought to interact with the C-terminus of N protein (*58, 60-63*). This interaction is required for maintaining packaging specificity in MHV (*64, 65*). The soluble CTD of M protein triggers RNA-independent phase separation when mixed with N protein (*35*), suggesting that M protein binding promotes a conformational change in N protein that leads to multivalent protein-protein interactions. Additionally, vRNPs in coronavirus virions appear to interact directly with the inner face of the virus membrane, with the circular ‘base’ of each vRNP cylinder proximal to the membrane (*43, 44, 57*). With these lines of evidence in mind, it seems likely that M protein binds ribonucleosomes, through the CTD and CTE of N, and tethers them to the viral membrane. Studies in MHV-infected cells have shown that N protein interacts with all coronavirus subgenomic RNAs, while M protein interacts only with full length genomic RNA (*63*). The interaction of M with the vRNP might therefore promote binding of specific sequences or structures in the SARS-CoV-2 genomic RNA that allows for exclusive packaging of the coronavirus genome. Further biochemical and genetic studies will be necessary to clarify the precise role of each protein in this process and to see if any specific sequences promote packaging specificity in SARS-CoV-2.

N protein is highly phosphorylated in infected cells, and numerous kinases have been implicated in this phosphorylation (*15, 16, 29, 30, 38*). Yaron et al. (*29*) recently provided evidence for sequential phosphorylation of N by SRPK, GSK3 and CK1 (Fig. 5A). Our results are consistent with this model. Phosphorylation of N is required for transcription of subgenomic RNAs that encode the four structural proteins (*16, 30*). We and others have previously shown that phosphorylated N forms liquid-like biomolecular condensates in the presence of viral RNA (*35, 38*). These condensates likely form at the RTC in infected cells and serve in part as a compartment to concentrate and protect viral replication and transcriptional machinery. N protein function at the RTC is also likely to depend, at least in part, on an interaction with Nsp3 on double membrane vesicles (*11, 32-34*). Our current results show that phosphorylated N protein cannot form ribonucleosomes, and instead forms elongated, heterogeneous ribonucleoprotein structures when mixed with longer viral RNA. These heterogeneous structures might serve as the foundation of the condensate compartment at the RTC and maintain RNA in an uncompacted state that is necessary for transcription of structural protein RNAs.

Chemical inhibition of SRPK with the FDA-approved drug Alectinib severely reduces replication of SARS-CoV-2 in multiple cell types (*29*). Additionally, inhibition of GSK3 with lithium reduces coronavirus replication in cultured cells, and analysis of clinical data of patients taking lithium revealed a ∼50% reduction in COVID-19 infection compared to those not on lithium (*66*). Thus, inhibition of N protein phosphorylation represents a promising target for therapeutic intervention that has the potential to reduce mortality in individuals infected with SARS-CoV-2 and could serve as an early treatment in the event of future coronavirus outbreaks.

## Materials and Methods

### N protein preparation

Wild-type and mutant N proteins were produced as described previously (*38*). Briefly, a codon-optimized synthetic DNA (Integrated DNA Technologies, IDT) was inserted into a pET28 expression vector by Gibson assembly, fused to DNA encoding an N-terminal 6xHis-SUMO tag. Mutant N proteins were generated by site-directed mutagenesis. N proteins were expressed in *E. coli* BL21 Star (Thermo #C601003), grown in TB-Kanamycin to OD 0.6, and induced with 0.4 mM IPTG. Cells were harvested, washed with PBS, snap frozen in LN_2_ and stored at -80ºC until use. Thawed cells were resuspended in buffer A (50 mM HEPES pH 7.5, 500 mM NaCl, 10% glycerol, 6 M urea) and lysed by sonication. The lysate was clarified by centrifugation and bound to Ni-NTA agarose beads (QIAGEN #30230) for 45 min at 4ºC. Ni-NTA beads were washed 3 times with 10 bed volumes of buffer A and eluted with buffer B (50 mM HEPES pH 7.5, 500 mM NaCl, 10% glycerol, 250 mM imidazole, 6M urea). The eluate was concentrated in centrifugal concentrators (Millipore Sigma #UFC803024), transferred to dialysis tubing (Spectrum Labs #132676), and renatured overnight by dialysis in buffer C (50 mM HEPES pH 7.5, 500 mM NaCl, 10% glycerol). Recombinant Ulp1 catalytic domain (purified separately from *E. coli*) was added to renatured protein to cleave the 6xHis-SUMO tag, and cleaved protein was injected onto a Superdex 200 10/300 size exclusion column equilibrated in Buffer C. Peak fractions were pooled, concentrated, frozen in LN_2_, and stored at -80ºC.

### RNA preparation

Sequences of all RNAs used in this study are provided in table S2. The template for in vitro transcription of 5’-600 RNA was a synthetic DNA (IDT), inserted by Gibson assembly into a pUC18 vector with a 5’ T7 promoter sequence. The 5’-600 insert, including the 5’ T7 sequence, was excised by EcoR1 digestion and purified by size exclusion chromatography on a Sephacryl 1000 column equilibrated in TE buffer (10 mM Tris pH 8, 1 mM EDTA). Peak fractions of the purified DNA insert were pooled and stored at -4°C.

Templates for all other long RNAs (5’-400, 5’-800, Nsp3, Nsp8/9, and Nsp10) were amplified by PCR of a plasmid containing the SARS-CoV-2 genome (a gift from Hiten Madhani, UCSF). All forward primers included a 5’ T7 promoter sequence. The SL8 and mSL8 templates were generated by PCR of synthetic DNA (IDT). The sequence for mutant SL8 (mSL8) was designed manually and checked for predicted secondary structure by RNAfold (http://rna.tbi.univie.ac.at/). PCR-amplified DNA was purified and concentrated by spin column (Zymo Research #D4004) before being used to generate RNA.

RNA synthesis was performed using the HiScribe T7 High Yield RNA synthesis kit (NEB #E2040S) according to the manufacturer’s protocol. Following incubation at 37°C for 3 h, in vitro synthesized RNA was purified and concentrated by spin column (Zymo Research #R1018). To promote formation of proper RNA secondary structure, all purified RNAs were heat denatured at 95°C for 2 min in a pre-heated metal heat block, and then removed from heat and allowed to cool slowly to room temperature over the course of ∼1 h. RNA concentration (A_260_) was quantified by nanodrop.

### Preparation of ribonucleoprotein complexes

The day before each experiment, N protein was dialyzed into reaction buffer (25 mM HEPES pH 7.5, 70 mM KCl) overnight. RNA was transcribed in vitro the day of analysis, heat-denatured and cooled slowly to allow for proper secondary structure. To assemble vRNP complexes, RNA was mixed with N protein (256 ng/μl RNA and 15 μM N, unless otherwise indicated) in a total volume of 10 μl and incubated for 10 min at 25°C. Samples containing stem-loop RNAs (SL4a, SL7, SL8, SL8m) were crosslinked by addition of 0.1% glutaraldehyde for 10 min at 25°C and then quenched with 100 mM Tris pH 7.5. Samples containing longer RNAs (5’-400, 5’-600, 5’-800, Nsp3, Nsp8/9, Nsp10) were not crosslinked. After assembly, vRNP complexes were analyzed as described below.

### RNA gel electrophoresis

After assembly (and crosslinking in the case of stem-loop RNAs), 10 μl vRNP mixtures were diluted 1:10 in dilution buffer (25 mM HEPES pH 7.5, 70 mM KCl, 10% glycerol). 2 μl of diluted vRNP mixtures was loaded onto a 5% polyacrylamide native TBE gel (Bio-Rad) and run at 125 V for 80 min at 4°C. 1 μl of the diluted samples was then denatured by addition of 4 M urea and Proteinase K (40 U/ml; New England Biolabs #P8107S), incubated for 5 min at 65°C, loaded onto a 6% polyacrylamide TBE-Urea Gel (Thermo Fisher), and run at 160 V for 50 min at room temperature. Gels were stained with SYBR Gold (Invitrogen) and imaged on a Typhoon FLA9500 Multimode imager set to detect Cy3.

### Mass photometry

Mass photometry experiments were performed using a OneMP instrument (Refeyn). A silicone gasket well sheet (Grace Bio-Labs) was placed on top of a microscope coverslip and positioned on the microscope stage. 10 μl reaction buffer (25 mM HEPES pH 7.5, 70 mM KCl) was first loaded into the well to focus the objective, after which 1 μl of vRNP complex sample was added to the reaction buffer, mixed, and measured immediately. Samples containing stem-loop RNA were diluted 1:10 before a second 1:10 dilution directly on the coverslip, while samples containing longer RNAs were only diluted 1:10 on the coverslip.

The mass photometer was calibrated with NativeMark™ Unstained Protein Standard (Thermo #LC0725). Mass photometry data were acquired with AcquireMP and analyzed with DiscoverMP software (Refeyn). Mass photometry data are shown as histograms of individual mass measurements. Peaks were fitted with Gaussian curves to determine the average molecular mass of the selected distributions. Each condition was independently measured at least twice.

### Turbidity analysis

Freshly prepared and renatured RNA was mixed with dialyzed N protein and incubated for 2 min at room temperature. Absorbance was measured at 260 nm and 340 nm using the Nanodrop Micro-UV/Vis Spectrophotometer. Turbidity was calculated by normalization of the 340 nm measurements to the absorbance value at 260 nm.

### Negative stain

For negative-stain EM, 2.5 μl of vRNP samples were applied to a glow discharged Cu grid covered by continuous carbon film and stained with 0.75% (w/v) uranyl formate. A Tecnai T12 microscope (ThermoFisher FEI Company) operated at 120 kV was employed to analyze these negatively stained grids. Micrographs were recorded at a nominal magnification of 52,000X using a Gatan Rio 16 camera, corresponding to a pixel size of 1.34 Å on the specimen. All images were processed using cryoSPARC. Micrographs were processed with Patch-Based CTF Estimation, and particles were picked using the blob picker followed by the template picker. Iterations of 2D classification generated final 2D averages.

### Polarization Anisotropy

Fluorescent RNA was ordered from IDT as a 10-nt degenerate sequence (random nucleotide at every position) with a 3’-FAM modification. N protein constructs were serially diluted in dialysis buffer, mixed with 10 nM fluorescent RNA and incubated at room temperature for 5 min. Fluorescence was measured on a K2 Multifrequency Fluorometer. RNA was excited with polarized light at 488 nm and emission was recorded at 520 nm. Data from three independent N protein titrations were fit to a one-site binding curve using GraphPad Prism to determine K_D_.

### Glycerol gradient centrifugation

Glycerol gradients were assembled as previously described, with slight modifications (*67*). Briefly, 10-40% glycerol gradients (dialysis buffer containing 10% or 40% glycerol) were poured and mixed with the Gradient Master (BioComp). For GraFix purification, fresh 0.1% glutaraldehyde was added to the 40% glycerol buffer prior to gradient assembly. vRNP samples (generally 75 μl of 15 μM N with 256 ng/μl RNA) were gently added on top of the assembled 5 ml gradients and samples were centrifuged in a prechilled Ti55 rotor at 35,000 rpm for 17 h. Gradient fractions were collected by puncturing the bottom of the tube with a butterfly needle and collecting two drops per well. For analysis by negative stain electron microscopy and mass photometry, peak fractions were combined and buffer exchanged using centrifugal concentrators (Millipore Sigma #UFC510024). Concentrated samples were then re-diluted 1:10 with dialysis buffer (0% glycerol) and re-concentrated. Samples were diluted and re-concentrated three times.

### Kinase reactions

Kinases were purchased from Promega (SRPK1: #VA7558, GSK-3β: #V1991, CK1ε: V4160). 1.25 μM N protein was incubated with 80 nM kinase for 30 min at 30ºC in kinase reaction buffer (25 mM HEPES pH 7.5, 35 mM KCl, 10 mM MgCl_2_, 1 mM DTT, 0.5 mM ATP and 0.001 mCi/mL ^32^P-γ-ATP). Reactions were quenched upon addition of SDS loading buffer for analysis by SDS-PAGE and autoradiography.

Phosphorylated protein for vRNP analysis was prepared in 90 μl reactions containing 16.5 μM N (WT or S188A + S206A) and 80 nM SRPK, GSK3, and CK1 in kinase reaction buffer. Reactions were incubated 30 min at 30ºC before addition of 5 mM EDTA. RNA was added to a final concentration of 256 ng/μl (which diluted N protein to a final concentration of 15 μM) and incubated at room temperature for 15 min. vRNP samples were analyzed by gradient centrifugation with crosslinker (GraFix) as described above.

## Acknowledgments

We thank the members of the Morgan laboratory for discussions and comments on the manuscript; and Conor Howard, Elise Muñoz, Hayden Saunders, and R. Das for discussions and technical assistance.

## Author contributions

C.R.C. and A.N.A. conceived the project and performed most experiments with guidance from D.O.M.; M.B. performed negative stain analysis with guidance from Y.C.; C.R.C. and D.O.M. wrote the paper with contributions from all authors.

## Data and materials availability

All data needed to evaluate the conclusions of the paper are included in the paper or supplementary materials.

## Figures and Tables

**Fig S1.**
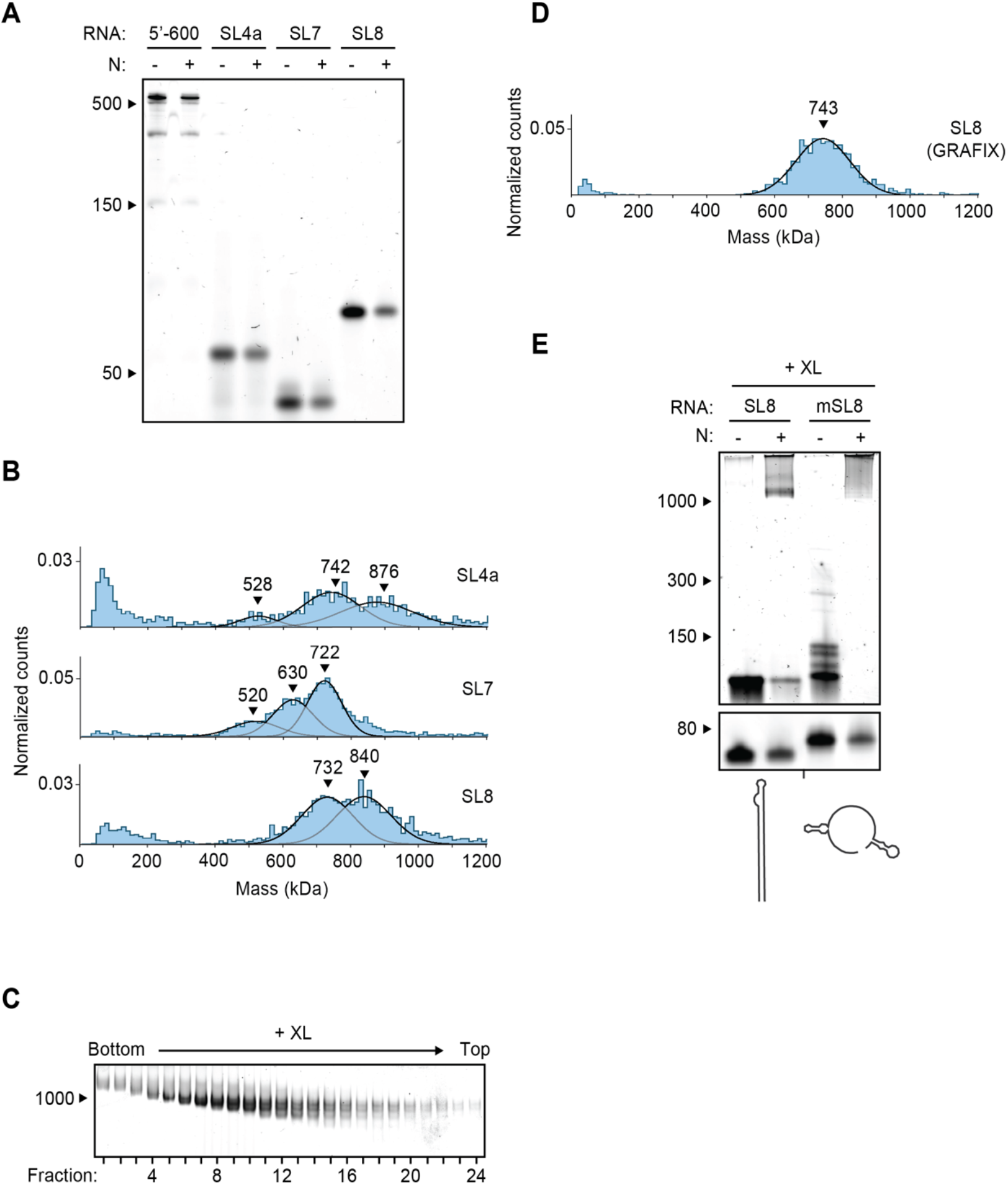
vRNP formation with stem-loop RNAs. (**A**) Denaturing gel electrophoresis of N protein mixed with indicated RNAs, related to Fig 2b. (**B**) Mass photometry analyses of crosslinked N protein complexes with indicated RNAs. Results with SL8 are reproduced in Fig. 2D. Representative of two independent experiments (table S1). (**C**) N protein in complex with SL8 RNA was separated by glycerol gradient centrifugation in the presence of crosslinker (GraFix) and analyzed by native gel electrophoresis. (**D**) Fractions 7 and 8 of GraFix-purified N-SL8 vRNPs (from C) were combined and analyzed by mass photometry. Representative of two independent experiments (table S1). (**E**) N protein was combined with SL8 RNA or mutant SL8 RNA (mSL8), crosslinked, and analyzed by native (top) and denaturing (bottom) gel electrophoresis. Predicted secondary structures are shown below. See table S2 for sequences.

**Fig S2.**
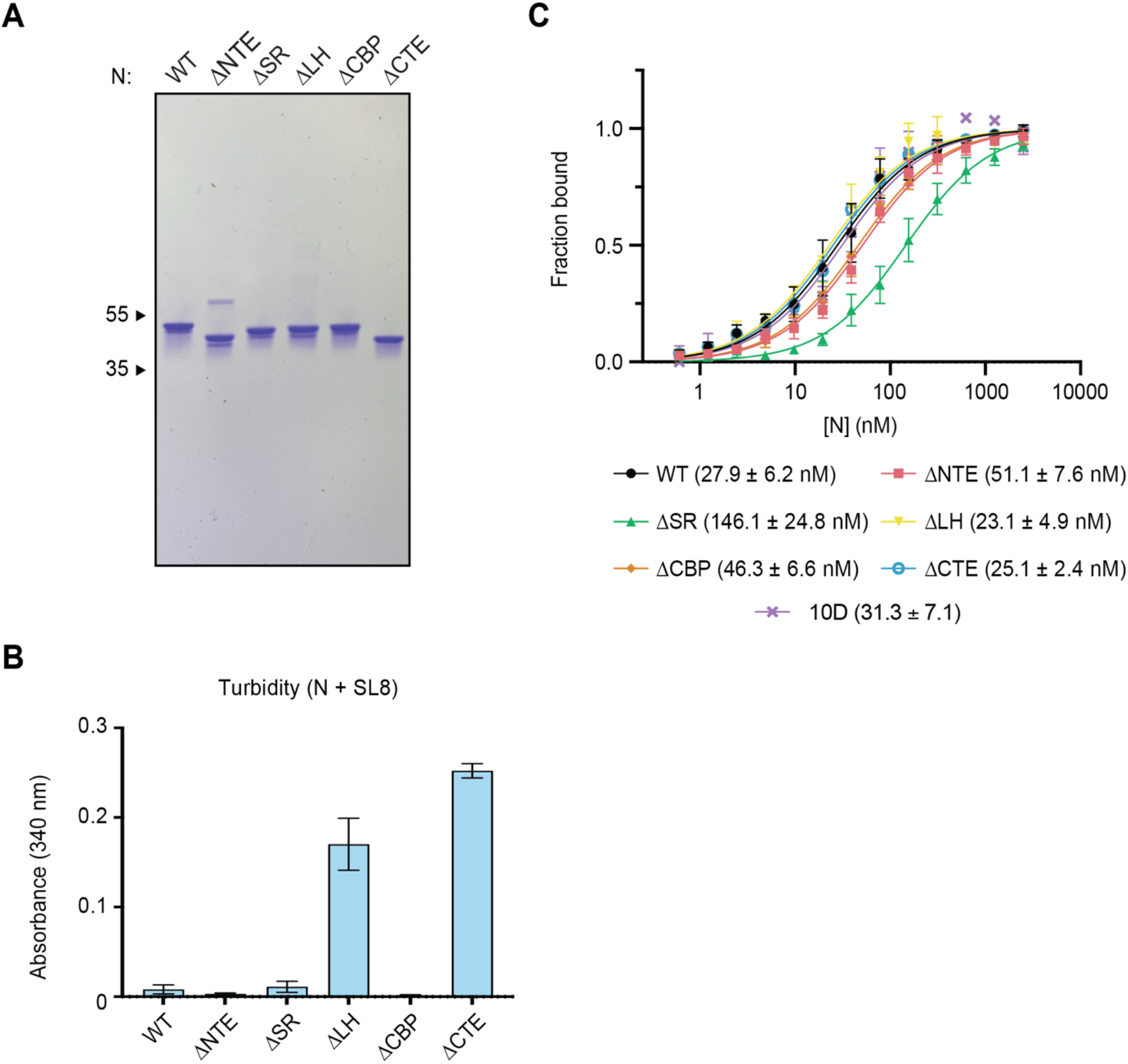
Analysis of N protein deletion mutants. (**A**) SDS-PAGE of N protein constructs used in this study, stained with Coomassie Blue. Molecular weight markers at left (kDa). **(B)** Absorbance at 340 nm was used to quantify the turbidity of wild-type and mutant N proteins mixed with SL8 RNA. All values are normalized to absorbance at 260 nm. (**C**) The indicated concentrations of N protein were incubated with 10 nM RNA (an entirely degenerate 10-nt RNA oligo with a 3’-FAM modification) and fluorescence anisotropy was measured. Data points reflect mean ± SEM of three independent experiments. K_D_ of each mutant is shown below.

**Fig S3.**
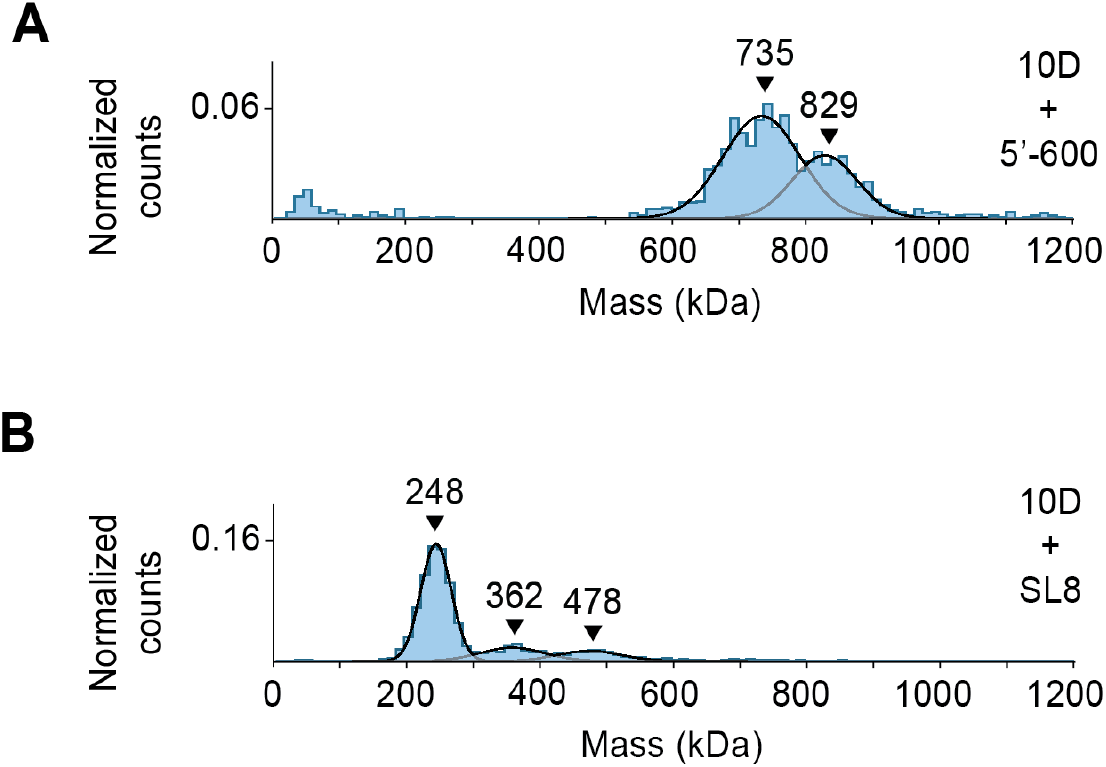
Analysis of complex formation by 10D mutant. Mass photometry analysis of GraFix-purified (**A**) fractions 7 + 8 of N protein in complex with 5’-600 RNA and (**B**) fractions 19 + 20 of N protein in complex with SL8 RNA. Representative of two independent experiments (table S1).

**Table S1.**
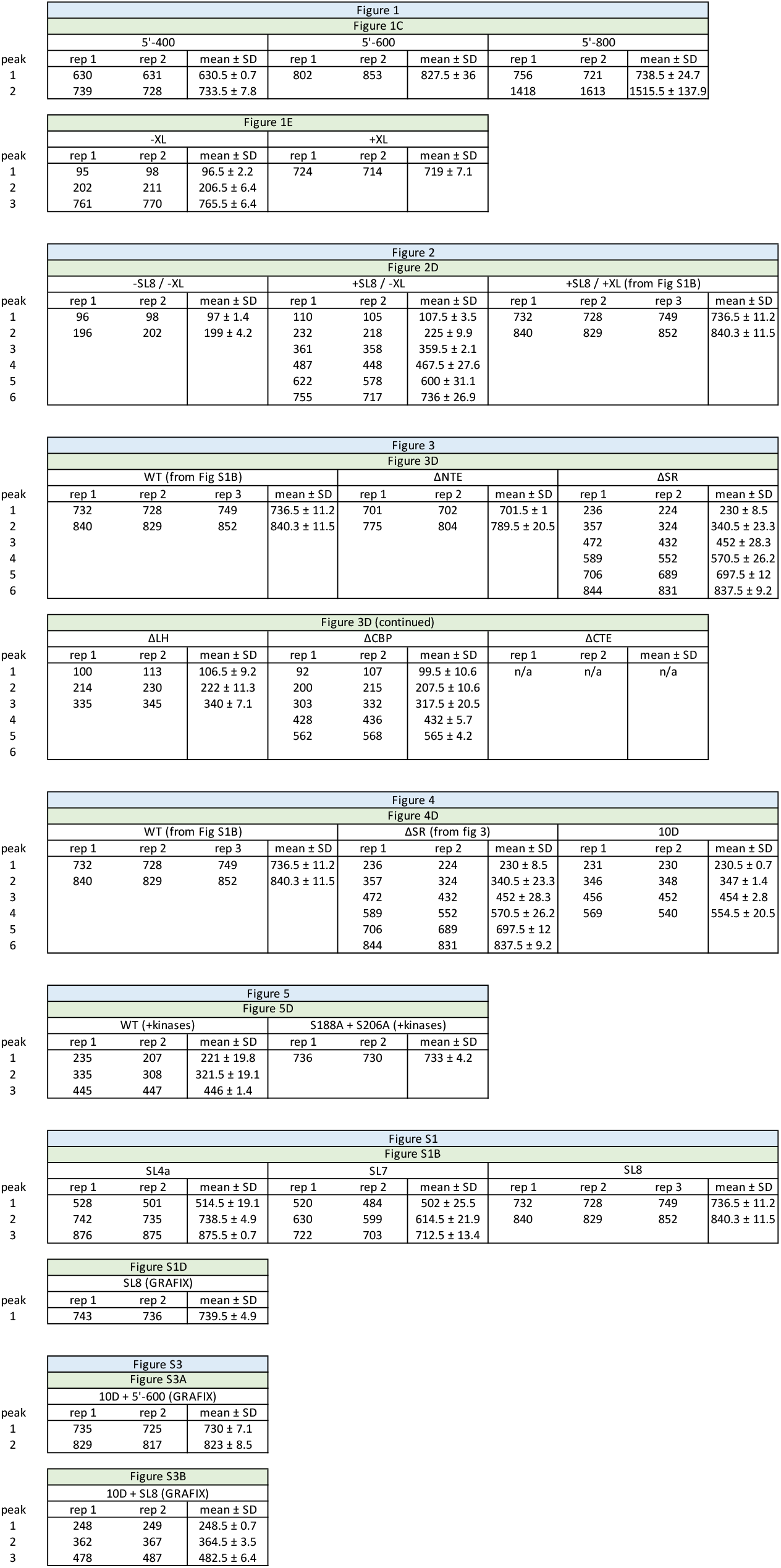
Summary of mass photometry results (kDa).

**Table S2.**
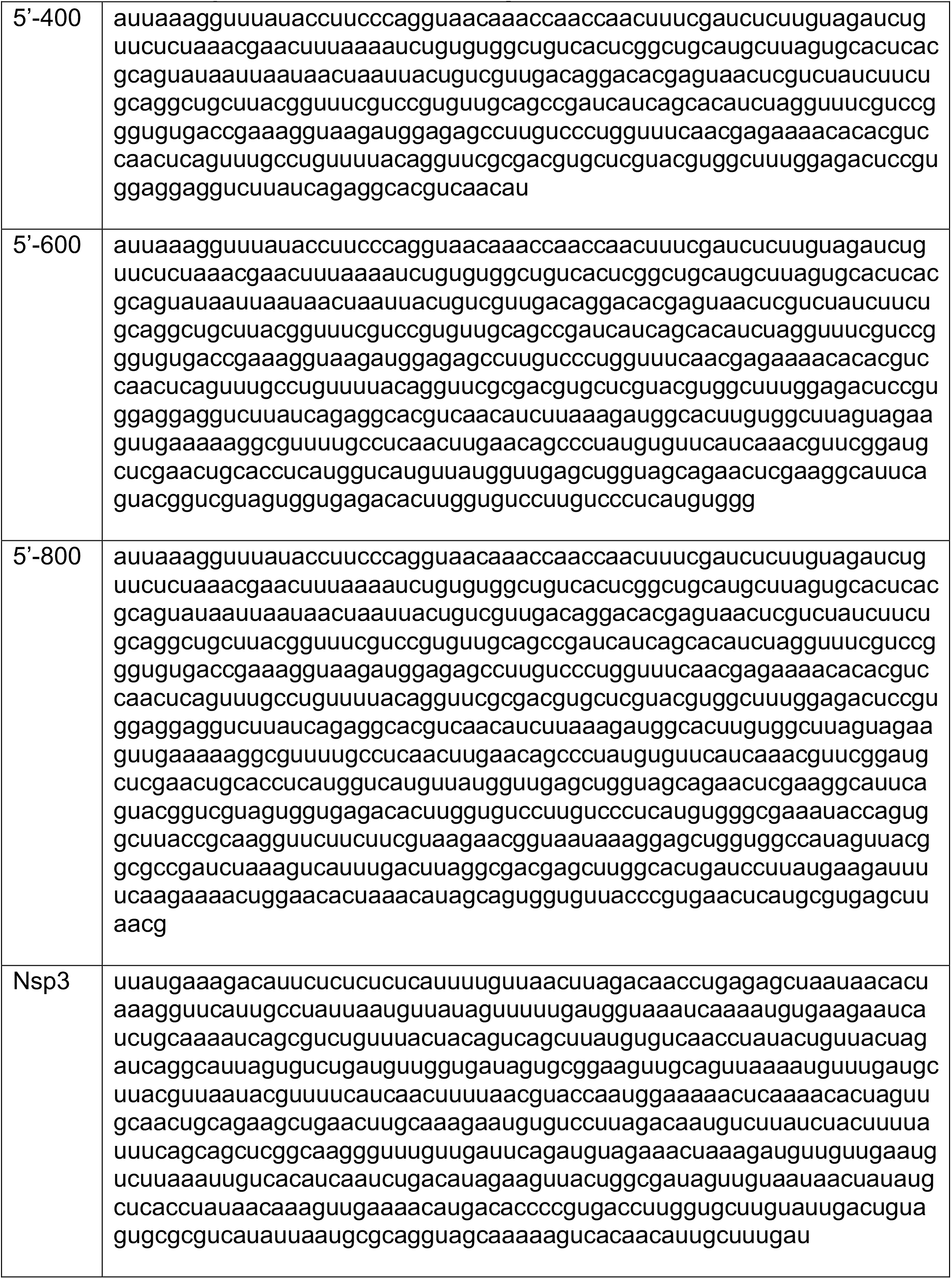
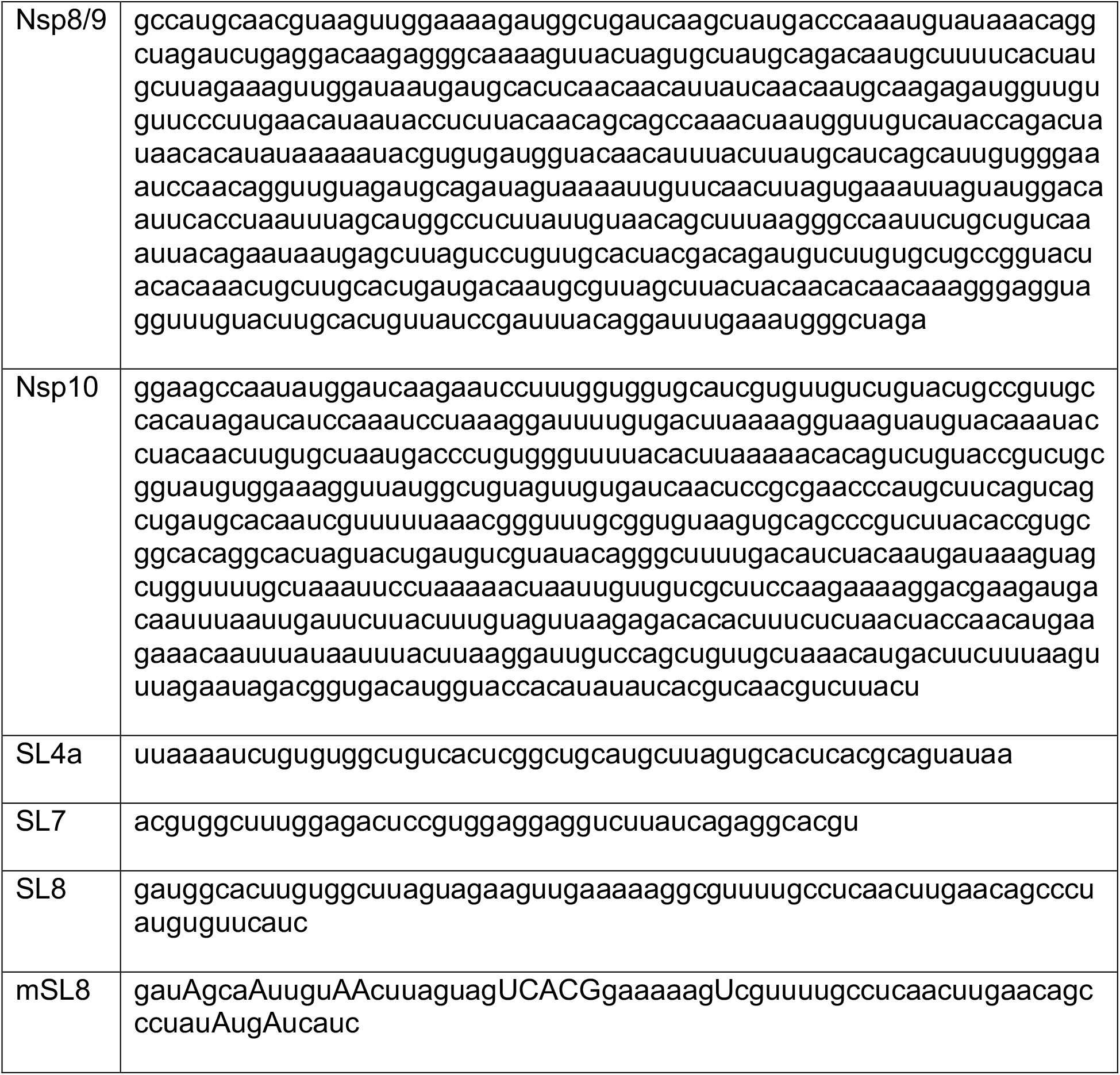
RNA sequences used in this study.

